# Pebbles as Dry Ice Replacement for Snap Freezing of Rodents Brains

**DOI:** 10.1101/100776

**Authors:** René Bernard, Larissa Mosch, Ulrich Dirnagl

## Abstract

Dry ice is commonly used as a cooling agent in the preclinical research environment. It permits rapid freezing of tissues and organs through liquid media. Laboratories depend on a constant supply of dry ice because it disintegrates in a matter of days. Here we tested whether commercially available pebble gravel could be used as dry ice replacement for snap freezing mouse brains in 2-methylbutane. We monitored 2-methylbutane temperature for one hour and tested different setups with the focus on creating a practical solution that can be used in every research laboratory. While gravel pebbles cannot replace dry ice in the laboratory entirely, they present a useful addition as readily available and reusable cooling agent.

## Introduction

Dry ice, which is frozen carbon dioxide, is next to liquid nitrogen, one of the most common cooling agents in the biomedical laboratory. It is used for intermediate term storage of samples either in the lab or for sample transport. It can also be used for cooling reagents down to -70°C. Dry ice usually comes in rectangular blocks of variable size. With mechanical force it can be crushed to increase cooling efficiency. While dry ice is versatile it has many shortcomings:

1. Dry ice needs to be produced and delivered from an external manufacturer.
2. Dry ice cannot be safely stored in -80°C freezers. Airtight freezers can cause dry ice to explode. It can also damage the freezers thermostat. (University of Rochester 2009)
3. Constant delivery and tedious forward planning is necessary to ensure on-demand-supply. This is challenging in a facility with multiple research groups depending on one central supply source for dry ice.
4. When in use, dry ice sublimates at room temperature, over one day a block of up to 4 kg dry ice can get lost if not stored in a well-insulated container or cooler.
5. Dry ice can burn the skin. Handling is only possible with insulated gloves to avoid frostbite.
6. Dry ice needs to be crushed with a hammer to increase surface contact for fast and efficient cooling.
7. Rooms in which dry ice is handled and stored need to be well ventilated to avoid CO_2_ buildup. When handling large amount of dry ice working under a hood is indicated.

Mainly due to the lack of alternatives dry ice remains one of the major cooling agents in research laboratories. In 2015 two different substitutes for dry ice (Ismalaj and Sackett 2015) were tested. The authors used either metallic beads or aquarium pebble gravel to cool down from room temperature a 4 ml ethanol/water (1:1) sample. They found that pebbles with 5 to 8 mm granular size were most effective in cooling 15 ml conical plastic tubes and that the rate of cooling was comparable to dry ice. The time course was limited to 5 min, however. Inspired by these results, we wanted to know whether pebbles can also be used for cooling down and keeping a liquid cold for a longer period. One prominent application ideally suited for testing pebbles in our lab is the cooling of 2-methylbutane, also known as isopentane. We use 2-methylbutane in a temperature range from -30°C to -45°C for snap freezing of freshly extracted mice brains prior to cryostat sectioning. In this context we were looking for an alternative to the constant supply and demand of dry ice. We developed and tested an approach that can be easily adapted by others with a focus on comparable reliability and cooling capacity as dry ice and optimized storage requirements for pebbles.

## Methods and Materials

A styrofoam box (22x16x18 cm) was used for storage of commercially available aquarium pebbles of 5-8 mm diameter (Wirbellosen Welt, Rödinghausen, Germany, http://www.wirbellose-online.de/Substrate/Bodengrund/Aquarienkies-Beige-5-8mm--10kg.html). One day prior to each experiment a pebble-containing box was stored in a -80°C freezer. Prior tests indicated that 4 hrs of freezer cooling is insufficient to reach -80°C pebble temperature. When the pebble-containing box was removed from the freezer, the experiment started immediately with the assessment of pebble temperature. Two different experimental setups were probed:

1. Pebbles were transferred into another, larger styrofoam box at room temperature. A 250 ml glass beaker (room temperature) was then centered and embedded in pebbles to maximize glass-pebble surface contact area.
2. The pebble-containing styrofoam box also contained a 250 ml glass beaker in the center, placed in the upright position, filled with paper towels and sealed shut with parafilm. (Figure 1A) Glass beaker and the surrounding pebbles were pre-chilled for 24 hr.

**Figure 1.**
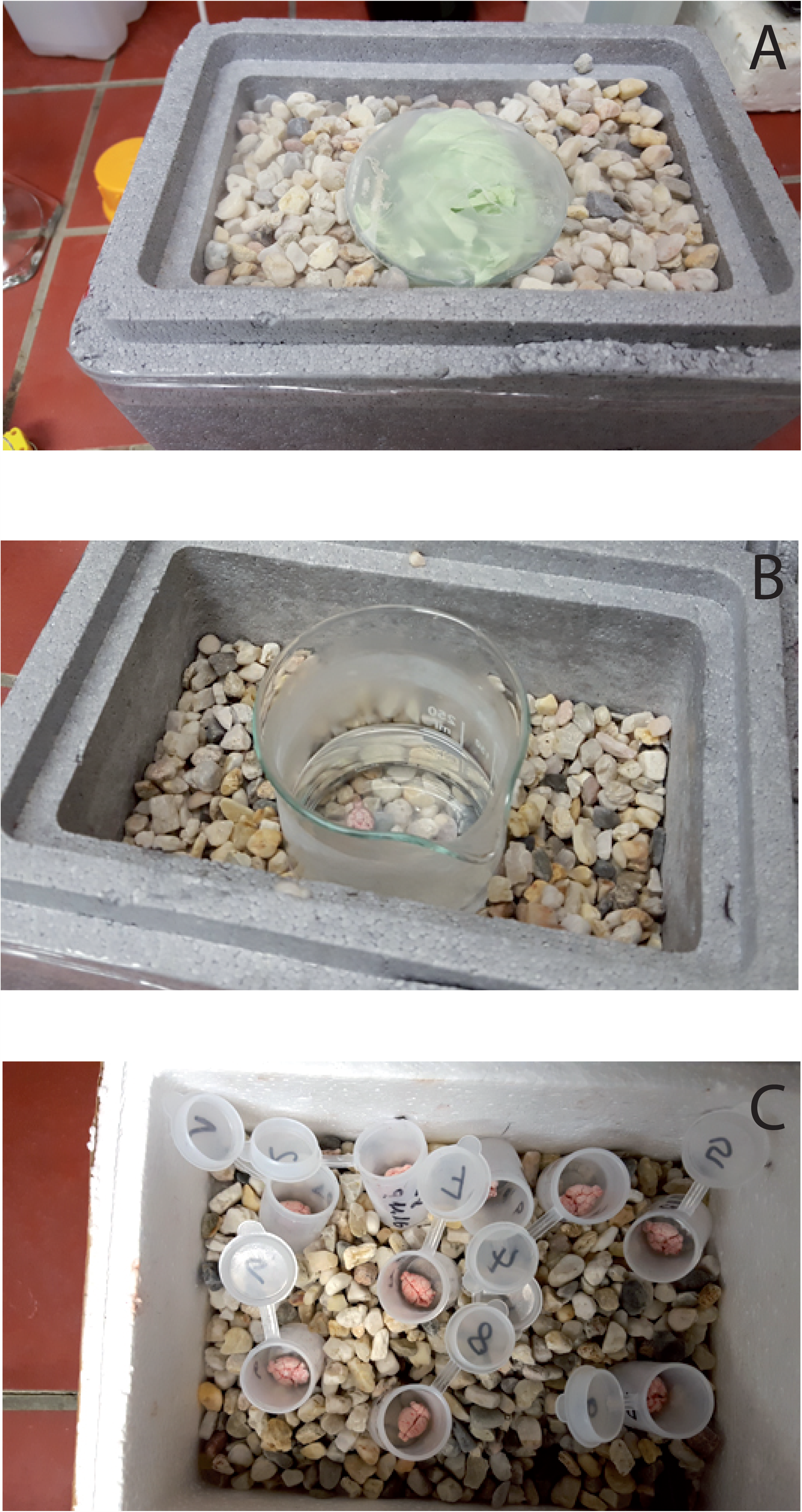
A) Glass beaker filled with paper towel and sealed with parafilm is packed in pebble-filled styrofoam box B) Glass beaker filled with 125 ml 2-methylbutane and pebbles filled up to liquid level C) Styrofoam storage box for frozen brains in plastic containers. This box uses the residue pebbles that were removed from the box shown in B.

In total, seven experiments were performed, three with setup 1, three using setup 2, and one control experimental with dry ice as cooling agent. Exactly 125 ml 2-methylbutane (Carl Roth GmbH, Karlsruhe, Germany) were filled in the pebble-surrounded glass beaker and temperature of 2-methylbutane was assessed every 5 min using a Voltcraft K101 digital thermometer connected to a 1 m long flexible temperature probe, with the tip placed in the center of the glass beaker, approximately 1 cm above the glass bottom. The aim was to assess the temperature of the liquid in an area where a brain would be located for snap freezing. At first cooling down time to approximately -30°C was measured. This was followed by the maintenance phase which lasted 60 min. In this phase temperature should be stable within the preferred interval for snap freezing. For three experiments we performed mouse brains extractions from 10-12 C57Bl6 mice and snap froze the brains in pebble-cooled 2-methylbutane to test the procedure under real conditions. Brain removal procedures were approved by the Landesamt für Gesundheit und Soziales, Berlin, and conducted in concordance with German animal protection act. Briefly, mice were deeply anesthetized by an intraperitoneal injection of a 4:1 mixture of ketamine 100 mg/ml and xylazine 0.8 mg/ml at a dose of 100 μl/g body weight) and then decapitated. Using surgical scissors and rongeur, skulls were carefully removed, the brains extracted and immediately placed in the prepared 2-methylbutane solution. After brains were solid frozen (<1 min) they were put into prelabeled round plastic containers which have been embedded in -80°C pebbles (Figure 1c). Brain extraction was performed continuously by a trained technician. All brains were in their containers in less than 60 min.

## Results

### Cooling with beaker at room temperature

When the room temperature-beaker filled with 2-methylbutane was embedded by the pebbles it took between 15-17 min until the target temperature of -30°C was reached (Figure 2a). In contrast, with dry ice this goal was achieved after approximately 6 min. When the cooling temperature was reached the temperature maintenance period started. For the first experiment (RT-beaker 1) we removed the beaker from the pebbles once temperature was going to drop below target range and we put it back into the pebbles when 2-methylbutane temperature was too warm for snap freezing. This approach mimicked the one of cooling 2-methylbutane with dry ice. During this pilot experiments with pebbles we noticed that the 2-methylbutane temperature did not change as rapidly as with dry ice and that the cooling capacity depends on by how much glas surface area was covered by the pebbles. For the following two experiments we covered only a portion of the glass beaker and had better sustainability of the 2-methylbutane temperature (Figure 2b). The average temperature of the pebbles at the end of maintenance period was -36.8 ± 2.9°C.

**Figure 2.**
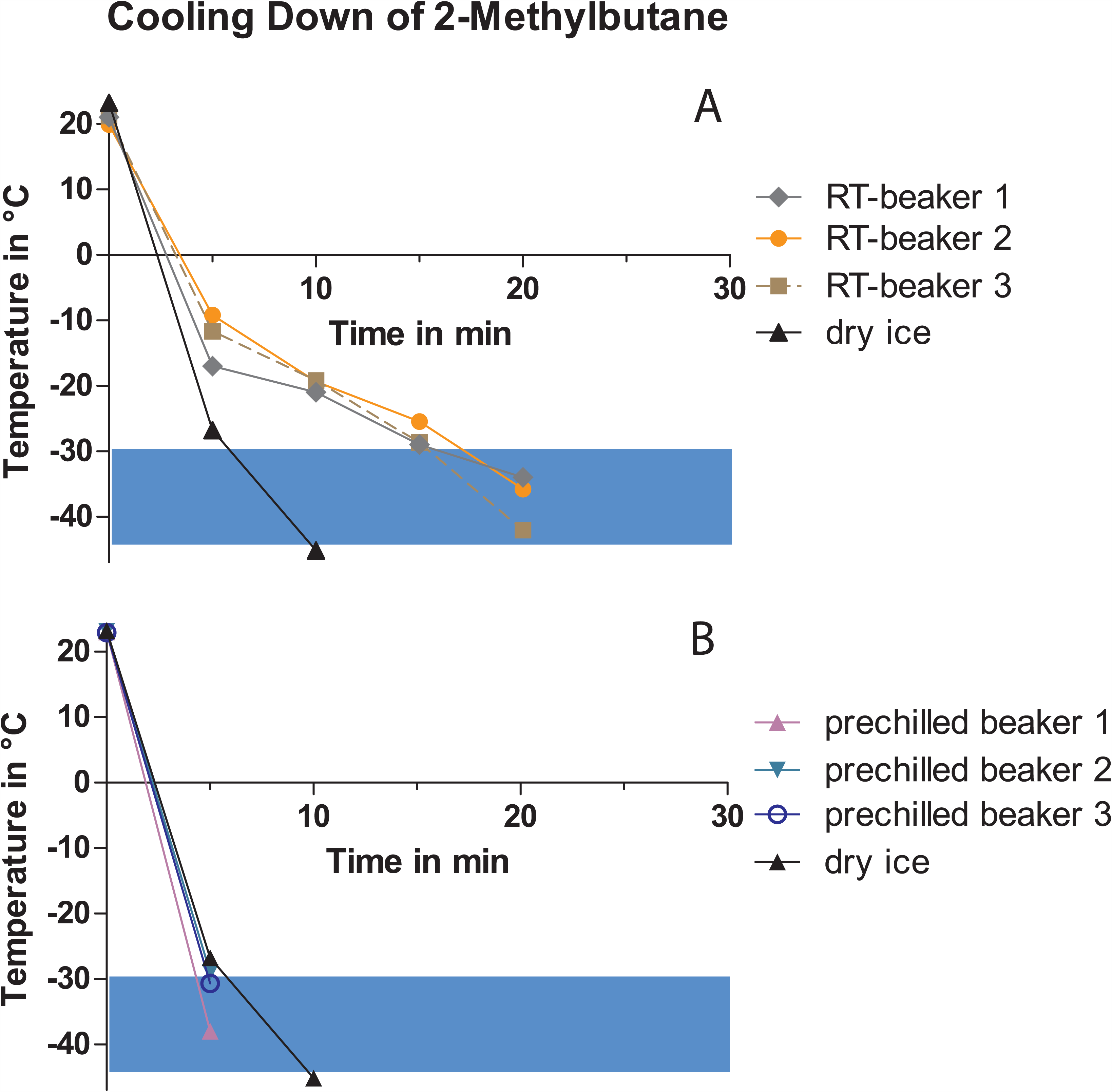
Cooling down of 2-methylbutane. Blue window indicates the target temperature zone. A) Time courses of cooling with pebbles and dry ice using glass beakers with room temperature at time point zero. B) Time courses of cooling with pebbles using a pre-chilled glass beaker. Dry ice time course is repeated from Fig.2A for comparison.

### Cooling with pre-chilled beaker

Placing the beaker in the pebbles and pre-cooling them together at -80°C affected the dynamics of 2-methylbutane cooling. As Figure 3a shows the preferred cooling temperature of -30°C was reached after 5 min. Under these condition cooling down velocity is comparable to that of dry ice. With pebbles covering the glass up to the rim, 2-methylbutane temperatures were below the target range. Therefore, we terminated this experiment after 45 min. For the next two experiments we placed the pebbles only to the liquid level of 2-methylbutane which limited the overall cooling capacity (Figure 1b). Temperature of 2-methyl butane was within the desired range for the entire maintenance period without human interference (Figure 3b). The average temperature of the pebbles at the end of maintenance period was -38.4 ± 1.2°C. For final experiment the pebbles was distributed as follows: 1650 gr for 2-methylbutane cooling and 650 gr for cooling box for brain containing plastic containers. Temperature of brains and containers were -20°C and -15°C respectively at the end of the final experiment (Figure 1c).

**Figure 3.**
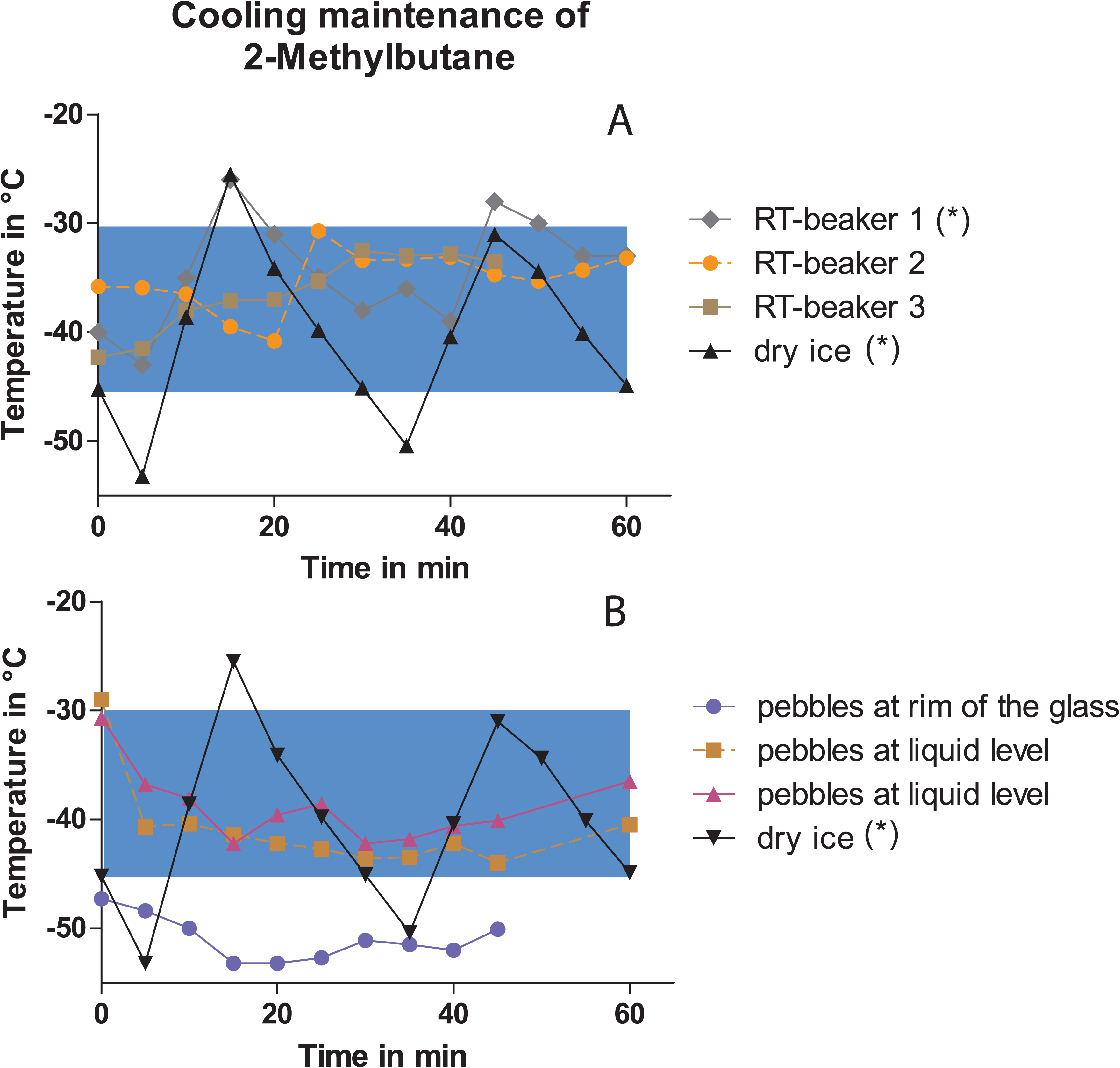
Cooling maintenance of 2-methylbutane. Blue window indicates the target temperature zone. (*) Glass beaker was removed from cooling source or reintroduced when 2-methylbutane temperature was outside the target range A) Time courses of cooling maintenance with pebbles and dry ice using glass beakers with room temperature at time point zero. B) Time courses of cooling maintenance with pebbles using a pre-chilled beaker. Dry ice time course is repeated from Fig.3A for comparison.

## Discussion

The present investigation showed that pebbles are capable to lower the temperature of 2-methylbutane to -30°C and to maintain this temperature for at least one hour. Cooling down of 2-methylbutane in a glass beaker that was kept overnight with pebbles at -80°C was faster than when the beaker had only room temperature. Prechilling might also increase the cooling capacity of pebbles beyond one hour and is therefore the preferred approach of the two. We set a one hour minimum because that is the maximum time needed for extracting brains from one experimental group, consisting of 10-12 mice, which requires immediate snap freezing. Our results indicate that pebbles are capable to cool 2-methylbutane sufficiently for this duration without the need to permanently check 2-methylbutane temperature.

### Freezing of biological tissue and organs

Tissue freezing is necessary to preserve morphology and to permit the detection of proteins or nucleic acids in qualitative and quantitative manner. It is crucial that freezing process is instantaneous because slow freezing causes ice crystal formation which introduces "swiss cheese" freezing artifacts in the cytoarchitecture. When a water-containing object is frozen rapidly, vitreous (amorphous) ice forms which does not expand when solidified (Scouten 2010). The liquid 2-methylbutane is an inert standard medium for tissue freezing. Dry ice is regularly used as cooling agent for lowering the temperature of 2-methylbutane to -30°C and below which avoids ice crystal formation in tissue. Liquid nitrogen, one of the coldest liquids available in biomedical laboratories, is unsuitable for freezing tissues blocks larger than 1 cubic centimeter. Liquid nitrogen creates a vapor barrier that causes freezing in an unpredictable manner due to gas layer build up through boiling which often causes cracks and brain tissue breakoffs (Scouten 2010).

Our results show that just like dry ice aquarium pebbles are capable of short term cooling 2- methylbutane in a range between -30°C and -45°C. Pebbles have different thermo-physical properties than dry ice. In opposite to pebbles, dry ice sublimates at room temperature creating a vapor barrier with sustainable low temperatures for many hours. However, gravel can cover a larger glass surface area than dry ice and absorb heat at a fast rate that is initially comparable to dry ice.

### Economical and ecological considerations

Dry ice cannot be produced in biomedical laboratories and is therefore not readily available. Depending on storage availabilities, delivery frequency and turnover, careful forward planning and organisation may be necessary. One drawback is the continuous decline in dry ice due to sublimation and that large amounts should not be stored in -80°C freezers. The cumulative demand for dry ice can be easily underestimated. The Department of Experimental Neurology of the Charité in Berlin centrally supplies dry ice to seven different research groups. In 2016 the annual consumption of dry ice in our department amounted to 1025 kg. The annual cost for dry ice are relatively low with 714 €. But compared to the cost of reusable pebbles of 3 € plus the electrical energy required for cooling, dry ice seems more expensive. Our annual usage of dry ice sets off 540 cubic meters carbon dioxide. Because dry ice is produced from gaseous carbon dioxide, the sublimation of dry ice into the atmosphere is considered emission-neural.

### Related applications and practical review

Many laboratories have limited storage capacity in -80°C freezers. One aspect of our setup was to minimize the spacial need for pebbles. We decided for a small styrofoam box for storage as container because it insulates pebbles against room temperature environment and without pebble transfer overall cooling efficiency is improved. This snap freezing method can also be used for any paraformaldehyde-perfused tissue after a cryoprotection step prior to storage at - 80°C (Rosene et al. 1986). The pebble-filled box can be also used for sorting samples that were kept at -80°C because Eppendorf and Falcon tubes glide perfectly within the pebbles. Frozen pebbles however, cannot replace dry ice entirely. There are applications that need long-term or intermittent cooling throughout an entire day for which dry ice is much better suited. In addition, dry ice provides more cooling capacity per gram, therefore dry ice is ideal for shipment of samples when the cool chain is to be maintained. The unavailability of dry ice can also endanger planned experiments. For instance, many experimental protocols require terminal blood collection together with organ collection. Under circumstance when dry ice is not available a protocol deviation would introduce unnecessary variability which exacerbates the comparability of measured parameters from blood or tissue samples. In such case, cooling with pebbles can act as experimental safeguard.

## Supplemental material

<u>Table S1</u> Excel file containing raw data from all conducted pebble and dry ice experiments with one experiment per sheet

